# Virtual tissue microstructure reconstruction across species using generative deep learning

**DOI:** 10.1101/2023.06.12.544541

**Authors:** Nicolás Bettancourt, Cristian Pérez, Valeria Candia, Pamela Guevara, Yannis Kalaidzidis, Marino Zerial, Fabián Segovia-Miranda, Hernán Morales-Navarrete

## Abstract

Analyzing tissue microstructure is essential for understanding complex biological systems in different species. Tissue functions largely depend on their intrinsic tissue architecture. Therefore, studying the three-dimensional (3D) microstructure of tissues, such as the liver, is particularly fascinating due to its conserved essential roles in metabolic processes and detoxification. Here, we present TiMiGNet, a novel deep learning approach for virtual 3D tissue microstructure reconstruction using Generative Adversarial Networks (GANs) and fluorescence microscopy. TiMiGNet overcomes challenges such as poor antibody penetration and time-intensive procedures by generating accurate, high-resolution predictions of tissue components across large volumes without the need of paired images as input. We applied TiMiGNet to analyze tissue microstructure in mouse and human liver tissue. TiMiGNet shows high performance in predicting structures like bile canaliculi, sinusoids, and Kupffer cell shapes from actin meshwork images. Remarkably, using TiMiGNet we were able to computationally reconstruct tissue structures that cannot be directly imaged due experimental limitations in deep dense tissues, a significant advancement in deep tissue imaging. Our open-source virtual prediction tool facilitates accessible and efficient multi-species tissue microstructure analysis, accommodating researchers with varying expertise levels. TiMiGNet’s simplicity and independence from paired images make it a versatile asset. Overall, our method represents a powerful and efficient approach for studying tissue microstructure, with far-reaching applications in diverse biological contexts and species.

## Introduction

The elucidation of tissue microstructure, encompassing the intricate arrangement and organization of cells, extracellular matrix, and other components within biological tissues, holds paramount significance in biological research(Mayer et al., 2012; Tanaka et al., 2017; Yadid et al., 2022; Yamaguchi et al., 2021). Of particular interest is the exploration of three-dimensional (3D) tissue microstructure, such as that observed in the liver, given its intricate organization and indispensable roles in metabolic, detoxification, and synthesis processes(Hoehme et al., 2010; Segovia-Miranda et al., 2019). The liver, comprising diverse cell types including hepatocytes, stellate cells, endothelial cells, and immune cells, exhibits a remarkably ordered arrangement(Morales-Navarrete et al., 2019a). The spatial distribution and interactions among these cells play a critical role in governing the liver’s proper functionality(Baratta et al., 2009; Bouwens et al., 1992). The hepatocytes together with the sinusoidal endothelial cells form complex tissue structures which optimizes the exchange of nutrients and bile secretion(Bouwens et al., 1992).

Unraveling the 3D microstructure of the liver and other tissues offers invaluable insights into tissue development, disease progression, and therapeutic responses(Segovia-Miranda et al., 2019). Perturbations in tissue microstructure often manifest during pathological conditions, including fibrosis, inflammation, and tumor growth(Abdelmalek, 2021). In-depth investigations of such alterations facilitate a deeper comprehension of disease mechanisms and the identification of potential targets for diagnostic and therapeutic interventions(Popa et al., 2021; Ray, 2020). Yet, capturing and reconstructing the 3D tissue microstructure accurately presents considerable challenges, even when using both traditional and advanced methodologies(Yoon et al, 2022). Reconstructing digital tissue models requires simultaneous imaging of multiple markers, such as antibodies, chimeric proteins, and small fluorescent molecules, across extensive volumes. The limitations of this endeavor encompass obstacles like inadequate antibody penetration, restrictions on the number of concurrently imaged fluorescent markers, and considerable acquisition time encompassing sample preparation and imaging procedures (Gigan et al, 2022, Yoon et al, 2022). Addressing these bottlenecks holds the key to advancing the field of 3D tissue modeling.

Over the past decade, deep learning has emerged as a groundbreaking technique in machine learning, showcasing its remarkable ability to unravel intricate patterns and representations from vast and complex datasets(LeCun et al., 2015). Deep learning has revolutionized the realm of artificial intelligence by eliminating the need for manual feature engineering. It employs artificial neural networks, inspired by the intricate connectivity of biological neural networks in the human brain, to automatically learn and discern intricate features directly from the data(Hallou et al., 2021; LeCun et al., 2015). By leveraging multiple layers of interconnected nodes, deep neural networks harness their hierarchical structure to progressively extract increasingly abstract and meaningful representations of the input data(Aloysius and Geetha, 2017). This learning technique has achieved unprecedented successes in critical tasks such as image classification, object detection, and image segmentation (Ravindran 2022). A distinctive area where deep learning’s prowess shines prominently is in the field of biology, particularly the analysis of microscopy images(Hallou et al., 2021; McQuin et al., 2018; Moen et al., 2019, Capec et al 2023). Deep learning models applied to microscopy image analysis have showcased their capability to automatically segment and classify cells, track their dynamic movements, and extract intricate features(Christiansen et al., 2018; Hallou et al., 2021; McQuin et al., 2018; Moen et al., 2019; Naert et al., 2021; Ounkomol et al., 2018; Xing et al., 2018). This transformative technology empowers researchers to investigate cellular dynamics, identify disease biomarkers, and expedite the discovery of novel therapeutics. Here, we propose an approach for virtual tissue microstructure reconstruction through the integration of Generative Adversarial Networks (GANs)(Goodfellow et al.) and fluorescence microscopy. By harnessing the strengths of GANs and leveraging the high-resolution imaging capabilities offered by fluorescence microscopy, we aim to provide a simple yet powerful tool to extract tissue micro-structural information from microscopy images of the actin meshwork. As a proof of principle, we reconstructed detailed and accurate reconstructions of the 3D tissue microstructure of liver samples of different species such as mouse and human. Our proposed approach holds high potential for enabling a deeper understanding of tissue microstructure in healthy and potentially diseased states.

## Results

### CNNs and GANs accurately predict several tissue structures in mouse liver tissue

Previous studies (Christiansen et al., 2018; Ounkomol et al., 2018) have shown the great potential of in-silico labeling by using deep learning models to predict fluorescent labels in unlabeled microscopy images. However, they are limited to cell culture systems. Here, we used deep learning models to generate in-silico labeling of 3D complex tissue microstructures. We used two well-established deep learning models, namely convolutional neural networks (CNNs) and Generative Adversarial Networks (GANs) ; and we compared their predictive power when applied to generate virtual images of various components of liver tissue microstructure based solely on high-resolution images of the actin mesh. These images were acquired using confocal microscopy. We used two network architectures: modified versions of UNet (Ronneberger et al., 2015) and CycleGAN (Zhu et al., 2017), termed TiMiPNet (Tissue Microstructure Predictor) and TiMiGNet (Tissue Microstructure Generator), respectively (refer to the Methods section for details). The TiMiPNet was trained using a dataset consisting of spatially registered pairs of 3D images of both the input and target structures in mouse liver tissue. Whereas images of the cell border (i.e. cortical actin mesh) served as the input, the output images included the BC network, sinusoids, or Kupffer cells (one target structure per trained network). Since, TiMiGNet (Figure 1a) does not require paired images, we computationally mixed the image pairs to simulate unpaired images and have a direct comparison with the models that require paired images such as TiMiPNet.

**Fig. 1:**
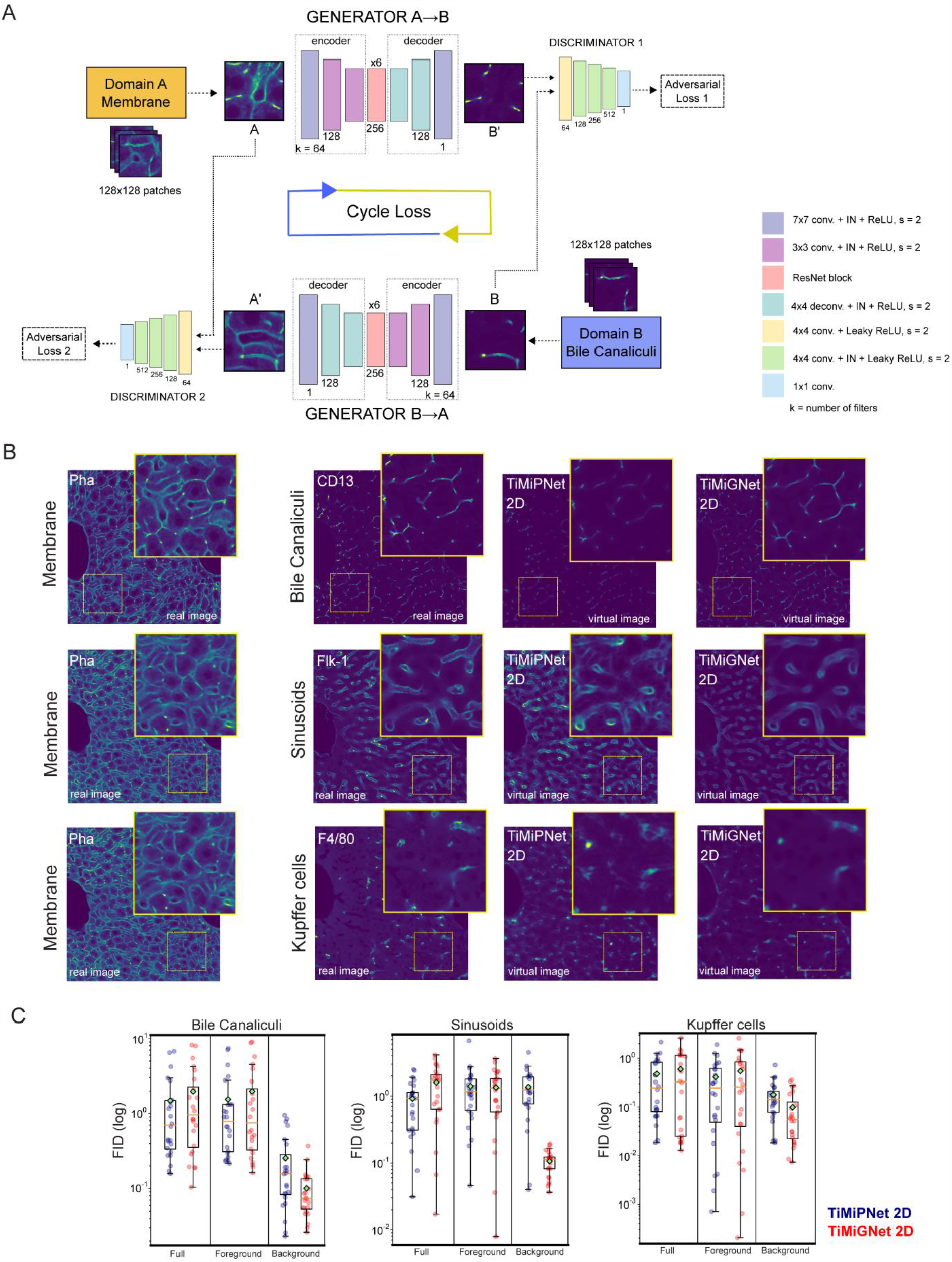
Deep CNN and GANs accurately predict liver tissue structures from cell border images. (a) Schematic representation of TiMiGNet. (b) 2D sections of 3D fluorescent images of the actin mesh (membranes), Bile Canaliculi, Sinusoids, and Kupffer cells (experimental images) as well as the corresponding predictions by Unet and CycleGAN in 2D. c) Quantification of the performance of the predictions of the BC network generated by the models. The test images were divided in 128x128x128 cubes and the metrics were estimated independently for each cube, i.e. each dot represents one image cube.

As shown in Figure 1-b and supplementary movies 1-6, both TiMiPNet and TiMiGNet generated outputs that closely resembled the ground truth images for the three predicted tissue structures, indicating their effectiveness in accurately predicting the different components of liver tissue microstructure from the actin mesh images, even in the absence of paired images (i.e. TiMiGNet). Next, we quantitatively evaluated the predictions of our models by comparing them with ground truth images acquired experimentally (Figure 1-b). Figure 1-c shows the Fréchet Inception Distance (FID) (Heusel et al., 2017), a widely used measure that assesses the likeness between the images produced by a generative model and the real images (see Methods for details), for TiMiPNet and TiMiGNet. We calculated the FID in the context of the entire images and by separately considering the background and foreground regions (Supplementary Figure 1a). Whereas lower values show good predictive power, high ones indicate larger difference between the prediction and the ground truth. As shown in Figure 1-c, both TiMiPNet and TiMiGNet show similar predictive levels for all structures (i.e. BC, Sinusoids and Kupffer cells).

To ensure a more comprehensive evaluation of our predictions, we conducted extra extensive tests specifically targeting the accuracy of predicting the BC, also comparing the results of both models in 2D and 3D. For a rigorous quantitative assessment, we employed various well-known evaluation metrics (See methods and Supplementary Figure 1-b). These metrics enabled us to thoroughly analyze the performance of our predictions. Detailed information regarding these evaluations can be found in the Methods section and Supplementary Figure 1. Interestingly, our results showed no substantial differences between the 2D and 3D architectures, as depicted in Supplementary Figure 1b. However, the computational costs for training 3D models are much higher than the ones needed for 2D architectures, i.e. up to 3 times higher for similar results. It is noteworthy that the majority of errors were primarily concentrated near the large veins, as illustrated in Supplementary Figure 2, as. Sudden changes in the morphology of the structures (i.e. BC) could be the source of these errors. Remarkably, both the TiMiPNet and TiMiGNet architectures consistently exhibited similar performance across all evaluated structures, as demonstrated in Figure 1-c and Supplementary Figure 1-b. This indicates that both approaches are equally effective in predicting the components of liver tissue microstructure. However, it is worth mentioning that the major advantage of the TiMiGNet approach is its ability to generate accurate predictions without relying on paired images. This characteristic enhances its flexibility and practical applicability in various scenarios.

### TiMiGNet accurately predicts BC and sinusoidal networks for deep tissue reconstructions

Given the generality and high performance demonstrated by TiMiGNet in our initial experiments, we proceeded to test its capabilities in more challenging tasks. In particular, we aimed to investigate whether TiMiGNet could effectively predict the intricate structures of the BC and sinusoidal networks in the context of deep tissue imaging. Traditional 3D imaging methods face inherent limitations when it comes to imaging deep tissue regions beyond a depth of approximately 80 to 100 micrometers, especially when utilizing antibody markers for specific structures such as the BC and sinusoids. The limited penetrance of antibodies within the tissue hampers the visualization of these structures beyond the shallow surface layers (Figure 2a, Suppl. movie 7-10). We took advantage of the unique properties of small fluorescent molecules, such as Phalloidin, which stains the actin mesh of the cells. Phalloidin has the remarkable ability to penetrate several hundred microns into the tissue, surpassing the constraints of antibody staining. We acquired high-resolution images of the actin mesh at a depth of ∼240 microns, well beyond the typical imaging range achievable with antibody-based techniques (Figure 2-a). Using our pre-trained TiMiPNet and TiMiGNet models, we then we predicted both the BC and sinusoidal structures from these deep tissue images. Our approach yielded accurate predictions of these structures throughout the entire depth of the images, exceeding the limitations imposed by the restricted antibody staining penetration depth of 80 microns. This is visually depicted in Figure 2-a and Supplementary movies 7-10. Figure 2-b and Figure 2-c show the mean intensity along the tissue depth for the BC and sinusoids, respectively. Whereas the intensity values for the experimental images, suddenly decreased after ∼80 µm, indicating lack of signal; the values for the virtual predictions remained stable along the whole tissue sample, showing both structures (BC and sinusoids) in places where no experimental staining existed (Suppl. movies 7-10).

**Fig. 2:**
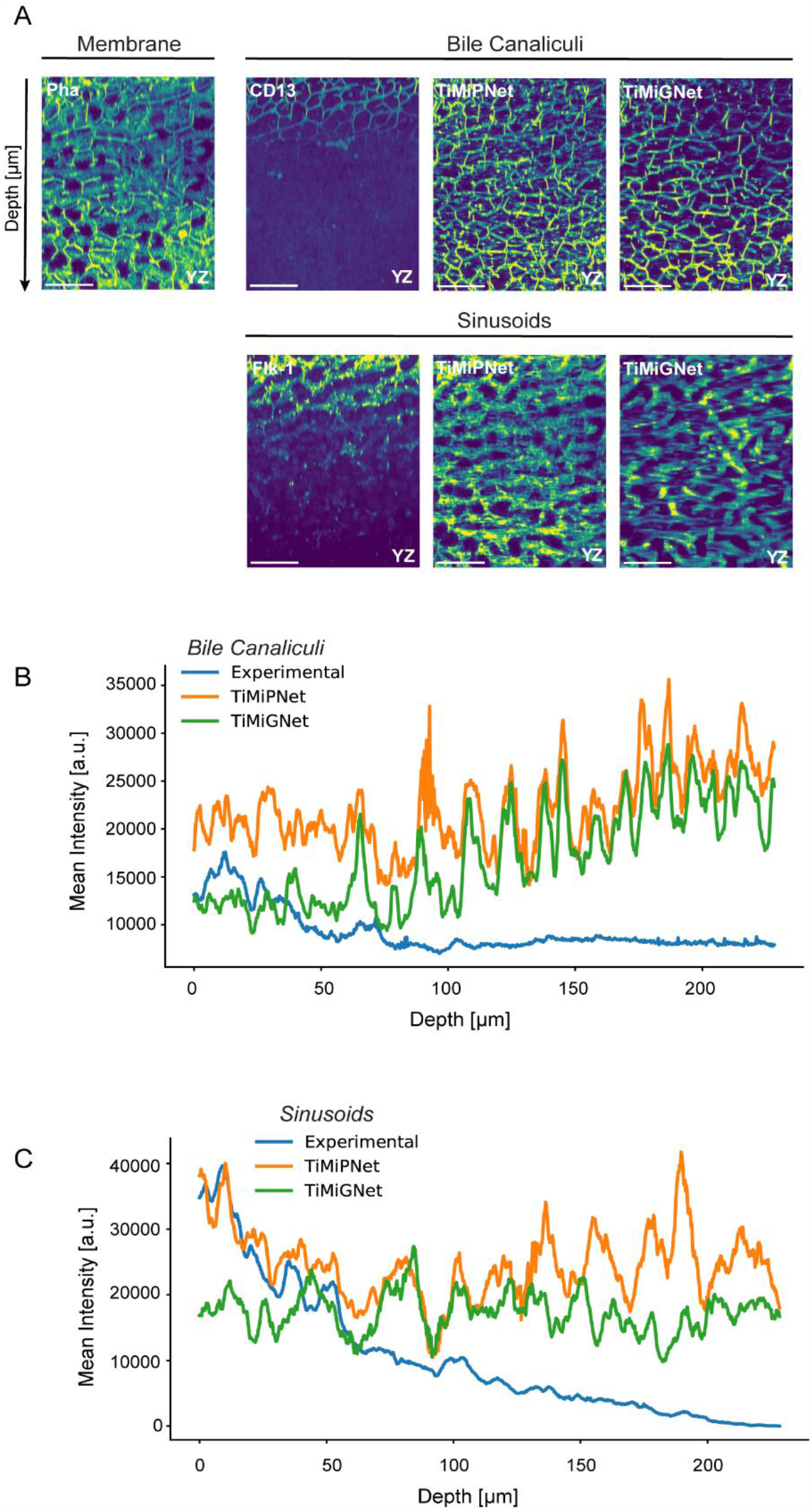
Deep tissue reconstructions using TiMiGNet. (a) 2D maximum projections of ∼20 µm of the axial sections of 3D fluorescent images of the actin mesh (membranes), Bile Canaliculi, Sinusoids (experimental images) as well as the predictions of TiMiGNet and TiMiGNet. (b) Quantification of the BC signal (mean intensity) along the tissue depth, in the corresponding images of panel a. (c) Quantification of the Sinusoids signal (mean intensity) along the tissue depth, in the corresponding images of panel a.

The successful application of TiMiGNet in predicting structures beyond the experimentally accessible depth of antibody staining without a need of paired ground truth images represents a great advancement in the field of deep tissue imaging. Our approach not only avoids the need for laborious, expensive and time-consuming antibody-based techniques but also offers a significant reduction in imaging costs. Moreover, by enabling the reconstruction of tissue structures that were previously inaccessible using conventional methods, our approach opens new avenues for exploring the intricate details of deep tissue microarchitecture. The ability to obtain deep tissue reconstructions using a readily available marker, such as Phalloidin, streamlines experimental procedures and expedites data acquisition. Furthermore, the successful prediction of structures beyond the limitations of antibody staining holds great potential for numerous research fields and clinical applications, revolutionizing our understanding of deep tissue biology and paving the way for new discoveries.

### Accurate prediction of tissue microstructure in human liver tissue

The general applicability of small dyes like phalloidin and DAPI enables the staining of tissues across various species, making them invaluable tools, especially when working with immunofluorescence, where challenges often arise due to the limited availability of antibodies that effectively function. To further push the boundaries of our approach, we sought to tackle an even more challenging task: predicting the bile canaliculi (BC) structures in human liver tissues. Overcoming this challenge required addressing the practical limitation of not being able to obtain paired images of the actin mesh and BC simultaneously.

For the human samples, we used images sourced from Segovia-Miranda et al. (2019). In their study, the authors encountered a challenge as only one antibody was found to effectively stain the BC network, and it requires an antigen retrieval protocol for optimal performance. However, this protocol disrupts the actin mesh, rendering phalloidin staining ineffective. Since simultaneous imaging of the actin mesh and BC in human tissue samples was deemed unfeasible, we trained our TiMiGNet model using separate sets of images representing the actin mesh and BC from different samples and patients. TiMiGNet exhibited a remarkable ability to generate visually convincing predictions of the BC in human liver tissue, as demonstrated in Figure 3b-middle and Supplementary movies 11-12. Moreover, we tested if the TiMiGNet trained in mouse tissue samples could also predict BC structures in human tissue. Surprisingly, TiMiGNet (trained in mouse tissue images) was also able to produce highly accurate predictions despite being originally trained on data from a different species (Figure 3b, bottom, and Suppl. movies 11-12). We quantitatively evaluate the predictions by comparing tissue parameters such as network radius and branch length estimated from experimental images and virtual predictions of the same patients but different tissue samples. Figures 3 b,c,d shows remarkably similar values for the predictions, despite the sample-to-sample variability previously shown in Segovia-Miranda et al., 2019. The values of the network radius and branch length were also evaluated along the liver lobule (CV-PV axis) to test spatial variability (Figures 3 c,d). It is worth noting that, even though the TiMiGNet model trained in mouse samples underestimates the BC radius in ∼10/20%, other morphological features such as the branch length are properly predicted. The fact that the network trained on mouse data could successfully generalize its predictions to human tissue underscores the remarkable conservation of BC structures across species. These findings provide valuable insights into the structural similarities and functional significance of BC in both mouse and human liver biology. The successful application of TiMiGNet approach in predicting BC structures in the absence of paired images highlights the adaptability and robustness of our method. Moreover, it suggests that the underlying architecture and organization of BC are highly conserved across different species.

**Fig. 3:**
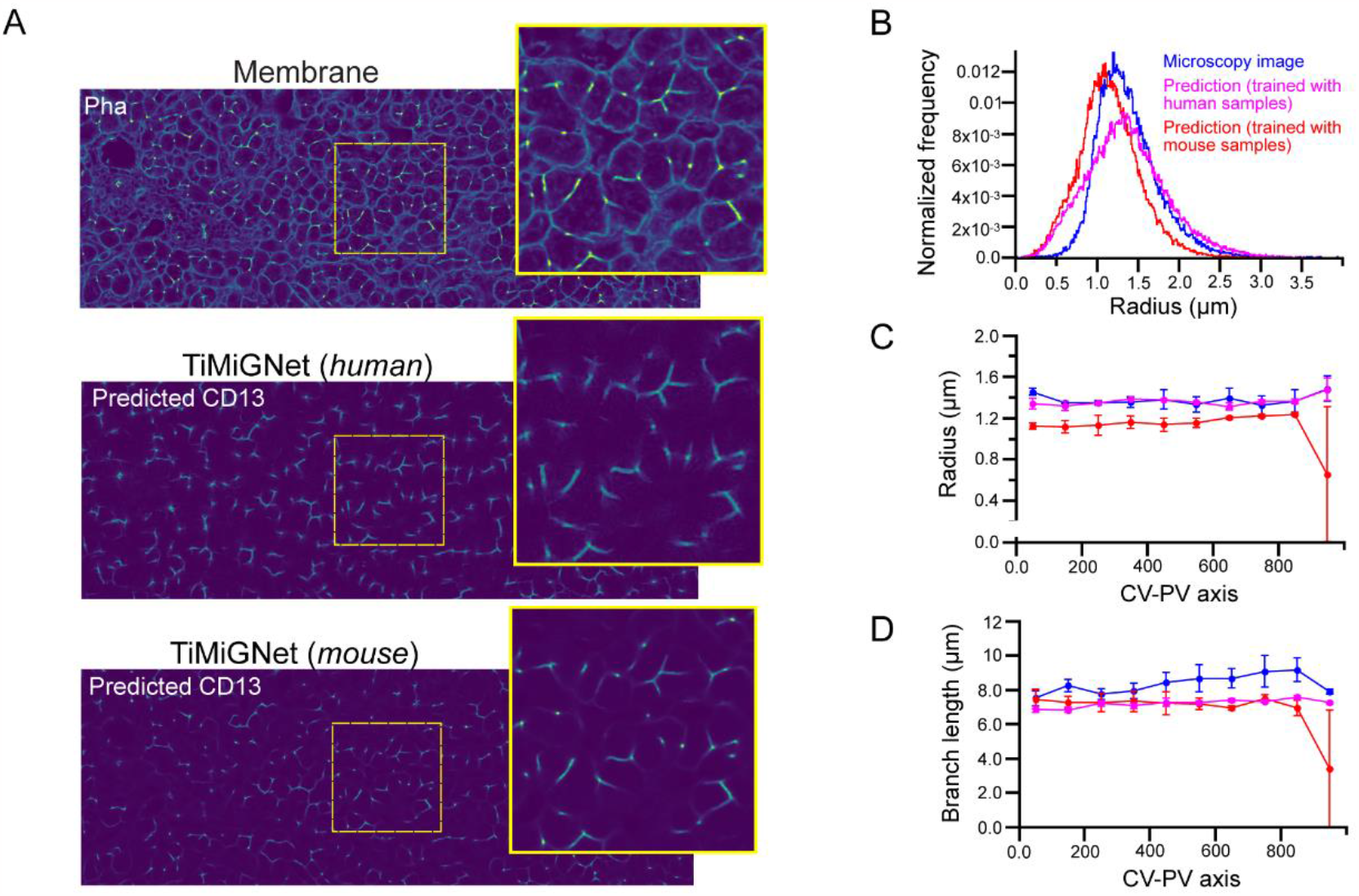
Prediction of tissue microstructure in human liver tissue using TiMiGNet. b) 2D sections of 3D fluorescent images of the actin mesh (membranes) and Bile Canaliculi (experimental images) of human liver tissue together with the corresponding predictions of the TiMiGNet 2D model trained in mouse and human tissue images. c-d) Quantification of morphological BC parameters for experimental images and TiMiGNet predictions: radius distribution(b), mean radius (c) and mean branch length (d) along the CV-PV axis.

## Discussion

Deep-tissue imaging and 3D reconstruction play pivotal roles in advancing our understanding of tissue architecture in both homeostasis and disease conditions. They enable the identification of different tissue characteristics, such as aberrant cell distribution, tissue remodeling, and the formation of disease-specific microenvironments. Current methods often suffer from limitations (poor antibody penetration, restrictions on fluorescent markers), require significant expertise, and can be time-consuming as they typically involve simultaneous imaging of multiple markers across large volumes (Gigan S. et al., 2022). Previously, we used CNNs to predict tissue structures by learning features embedded within single-marker images(Morales-Navarrete et al., 2019b). In particular, our deep learning framework showed remarkable accuracy for the prediction of the bile canaliculi (BC) and sinusoidal networks from images of the actin meshwork of liver tissue. However, this approach has several limitations: i) requiring a pair of images of the BC/Sinusoids and the actin mesh, which is not always technically possible, ii) using a 2D approach to predict 3D structures could cause loss of data information, iii) is limited to healthy mouse liver tissue. Here, we overcame these issues and generalized the approach by using Generative Adversarial Network, TiMiGNet. We showed that our proposed methodology has the capability to produce precise and high-resolution predictions of various tissue components, such as bile canaliculus, sinusoids, and Kupffer cell shapes, based on cell border images (actin meshwork), thereby facilitating efficient and dependable analysis. The integration of TiMiGNet with fluorescence microscopy allowed us to predict tissue structures in scenarios where paired images of ground truth were not attainable. For instance, we demonstrated the potential of using TiMiGNet in predicting structures beyond the experimentally accessible depth of antibody staining, representing a significant advancement in the field of deep tissue imaging. Moreover, we showed that TiMiGNet facilitated multi-species analysis (mouse, human). This makes TiMiGNet a powerful and versatile tool for researchers and practitioners in various domains of biology and medicine.

The utilization of a simple marker, such as Phalloidin, in our method further enhances the practicality and broadens the applicability of our method. Unlike the complex and time-consuming procedures required for simultaneous imaging of multiple markers, our approach relies on a single marker, making it more efficient and cost-effective. This simplicity enables researchers to obtain deep tissue reconstructions with ease, providing a valuable tool for investigating tissue microstructure. Moreover, our method could potentially even be applied in the absence of specific markers, if manually annotated structures are provided as input. This flexibility would allow to utilize our approach in situations where specific markers may not be available or suitable.

The ability to predict BC structures in human liver tissue using TiMiGNet, despite the inherent challenges and limitations, represents a significant advance. This achievement not only could help us to expand our understanding of liver tissue microarchitecture in the context of human biology but also opens up new avenues for investigating the role of BC in various liver diseases and clinical applications. Our results demonstrate the power of leveraging computational methods and deep learning techniques to overcome experimental constraints and provide valuable insights into complex biological systems. The successful translation of our approach from mouse to human liver tissue holds great promise for advancing our understanding of liver biology and facilitating the development of novel diagnostic and therapeutic strategies for liver-related disorders.

Our study presents a novel approach for analyzing tissue microstructure using well-established yet powerful deep learning models and demonstrates its effectiveness in predicting structures beyond the limitations of conventional methods. The simplicity and versatility of our method, particularly with the use of a single marker or minimal annotations, make it a valuable tool for research in various systems. TiMiGNet is made available to the community as open-source software through our GitHub repository (http://github.com/hernanmorales-navarrete/TiMiGNet) and includes the data sets used for training and testing via a Zenodo link. Its straightforward yet efficient design allows for seamless adaptation to diverse applications, ensuring its versatility for various purposes. Future improvements, such as expanding the training dataset and refining the network architecture, have the potential to further enhance the accuracy and reliability of our predictions, ultimately advancing our understanding of tissue microarchitecture and its implications in biological systems.

## Methods

### Animals

Adult C57BL/6J mice (8-10 weeks old) were obtained from the animal facility (Centro Regional de Estudios Avanzados para la Vida (CREAV)) at the Universidad de Concepción. The animals were maintained in strict pathogen-free conditions and received *ad libitum* feeding. All procedures performed were approved by the vice rectory of ethics and biosecurity committee from the investigation and development of Universidad de Concepción. Z-stack images of NAFLD human liver samples were obtained from(Segovia-Miranda et al., 2019).

### Mice liver collection and immunostaining

Mice livers were fixed through intracardiac perfusion with 4% paraformaldehyde 0.1% Tween-20/PBS and post-fixed overnight with the same solution at room temperature. 100 µm thick liver sections were obtained with a vibratome. Immunolabeling and optical clearing were performed as described previously(Morales-Navarrete et al., 2019a).

### Imaging

Liver samples were imaged (0.3 µm voxel size) in an inverted multiphoton laser-scanning microscope (Zeiss LSM 780) using a 40x1.2 numerical aperture multi-Immersion objective (Zeiss). DAPI was excited at 780 nm using a Chameleon Ti-Sapphire 2-photon laser. Alexa Fluor 488, 555 and 647 were excited with 488, 561 and 633 laser lines and detected with Gallium arsenide phosphide (GaAsp) detectors.

### Image pre-processing

The different components of liver tissues (BC, sinusoids and cortical mesh) were imaged with high-resolution (voxel size 0.3 x 0.3 x 0.3 µm) fluorescent image stacks (80/100µm depth). To cover the entire CV-PV axes, 2x1 tiles were stitched using the image stitching plug-in of Fiji(Preibisch et al., 2009). The 3D images were first denoised using the PURE-LET algorithm(Luisier et al., 2010) with the maximum number of cycles. Then, a background and shading correction was performed using the tool BaSiC(Peng et al., 2017) along the stack. Finally, all channels were aligned to the actin mesh channels using the function Correct 3D Drift from Fiji.

### TiMiGNet Architecture

The model consists of two Generators and two Discriminators. Generator A to B (G-AB) takes an image (real) of class A as input and generates an image (fake) of class B as output. On the other hand, Generator B to A (G-BA) takes an image (real) of class B as input and generates an image (fake) of class A as output. Discriminator A (D-A) classifies images generated by G-BA as real or fake, whilst Discriminator B (D-B) classifies images generated by G-AB as real or fake. The objective is to train both generators and both discriminators. This process will, eventually, allow the generators to create realistic enough fake images and deceive the discriminators.

#### Generator architecture

The architecture for models trained with patches of size 128x128 pixels: c7s1-64, d128, d256, R256 (x6), u128, u64, c7s1-1. The architecture for models trained with patches of size 256x256 pixels: c7s1-64, d128, d256, R256 (x9), u128, u64, c7s1-1. Where: c7s1-k is a convolution of k filters of size 7x7 and stride 1, followed by an Instance Normalization (IN) layer and a ReLU layer, dk is a convolution of k filters of size 3x3 with stride 2, followed by an IN layer and a ReLU layer, Rk is a residual block with two convolutions with equal number of filters of size 3x3, uk is a block of a transposed convolution with k filters of size 3x3 and stride 2, followed by an IN layer and a ReLU layer. Discriminator architecture: The architecture for both discriminators is: C64 - C128 - C256 - C512 - F1. Where: Ck is a block of a convolution of k filters of size 4x4, followed by an IN layer and a LeakyReLU layer, F1 is a convolution of 1 filter of size 4x4.

The network for the 3D model is essentially the same as the 2D model, with minor differences only: the 2D input and output layers were adapted to accept and produce 3D patches, respectively, and all 2D convolution layers were swapped for 3D convolution layers.

### Model Training

#### TiMiGNet 2D

Four images, two for each domain, were used to train the 2D models. Pixel values were normalized between -1 and 1. 2500 two-dimensional patches were extracted from each image, leading to 5000 patches per domain. Patches of size 128x128 pixels were used for the membrane/BC and the membrane/sinusoids models, whereas patches of size 256x256 pixels were used for the membrane/Kupffer model. All models were trained for 100 epochs using Adam optimizer with a learning-rate of 0.002, a beta value of 0.5, a batch-size of 1 and Mean Squared Error as the loss function.

#### TiMiGNet 3D

Four images, two for each domain, were used to train the 3D model. Pixel values were normalized between -1 and 1. For the membrane/BC model, 900 three-dimensional patches of size 64x64x64 pixels were extracted from each image, leading to 1800 patches per domain. The ninety percent of the patches were used for training and the remaining ten percent were used for validation. Hyperparameters for the 3D model were set to be equal to those of the 2D model, except for the number of epochs which was set to 300. This allowed us to stop the training early if no improvements were observed.

### Quality evaluation metrics

The test images were divided in 128x128x128 cubes and the metrics were estimated independently for each cube. We evaluated the predictive power of the models using an extensive set of well-establish metrics including:

Fechet Inception Distance (FID) [1], which measures the similarity between the generated and ground truth images based on features of the raw images calculated using the inception v3 model. FID is calculated by computing the Fréchet distance between two Gaussians fitted to feature representations of this model.

Mean Squared Error (MSE) [2], which measures the average squared difference between the pixel values of an image generated by a model and the pixel values of the ground truth image.

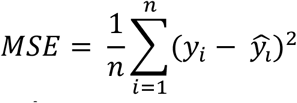

Where ŷ is the predicted image and ŷ is the ground truth image, *n* is the number of pixel/voxels in the images

Mean Squared Logarithmic Error (MSLE) [2] is similar to MSE but is applied to the natural logarithm of the generated and ground truth images. It is often used in tasks where the error distribution is expected to be logarithmic in nature.

Mean Absolute Error (MAE) [2] calculates the average absolute difference between the pixel values of the generated and ground truth images, making it less sensitive to outliers compared to MSE.

Root Mean Squared Error (RMSE) [2] is the square root of the MSE. It provides a measure of the standard deviation of the errors.

Peak Signal-to-Noise Ratio (PSNR) [3] is an expression for the ratio between the maximum pixel value of the ground truth image to the Mean Squared Error between the generated and ground truth images.

Structural Similarity Index Measure (SSIM) [3] is a metric used to assess the structural similarity between two images based on a perception-based model that takes into account image luminance, contrast, and structure.

Multi-scale Structural Similarity Index Measure (MS-SSIM) [4] is an extension of SSIM that considers multiple scales in the image. It provides a more comprehensive assessment of image quality by taking into account variations at different levels.

Cosine Similarity (COS) [5] is a measure used to determine the similarity between two vectors in a multi-dimensional space. It calculates the cosine of the angle between the vectors.

Correlation Coefficient (CoC) [5] is a statistical measure that quantifies the linear relationship between two sets of data points. For 2D images it can be used to assess how closely related or linearly associated are the pixel values of the generated and ground truth images.

## Author Contributions

H.M-N. conceived the project. H.M-N and F.S-M designed and supervised the project. H.M-N, F.S-M, Y.K and M.Z. developed the conceptual bases of the project. V.C., C.P., and F.S-M. performed the immunofluorescence, optical clearing and imaging. N.B and H.M-N. implemented the deep convolutional neural networks and analysis pipelines. N.B., and H.M-N and C.P. performed the data analysis and interpretation of the results. H.M-N., F.S-M., N.B and P.G. wrote the manuscript.

## Conflict of interest

Authors declare no conflict of interest.

### Acknowledgments

We would like to thank the following Facilities from Universidad de Concepción for their support: Centro de Microscopía Avanzada (CMA BIO-BIO) and Centro Regional de Estudios por la Vida (CREAV).

## Data and code availability

The TiMiGNet open-source code along with example data is available from http://github.com/hernanmorales-navarrete/TiMiGNet. All the image data used for training and test is available at *zenodo_link_upon_publication*.

## Figures

**Supp Fig. 1:**
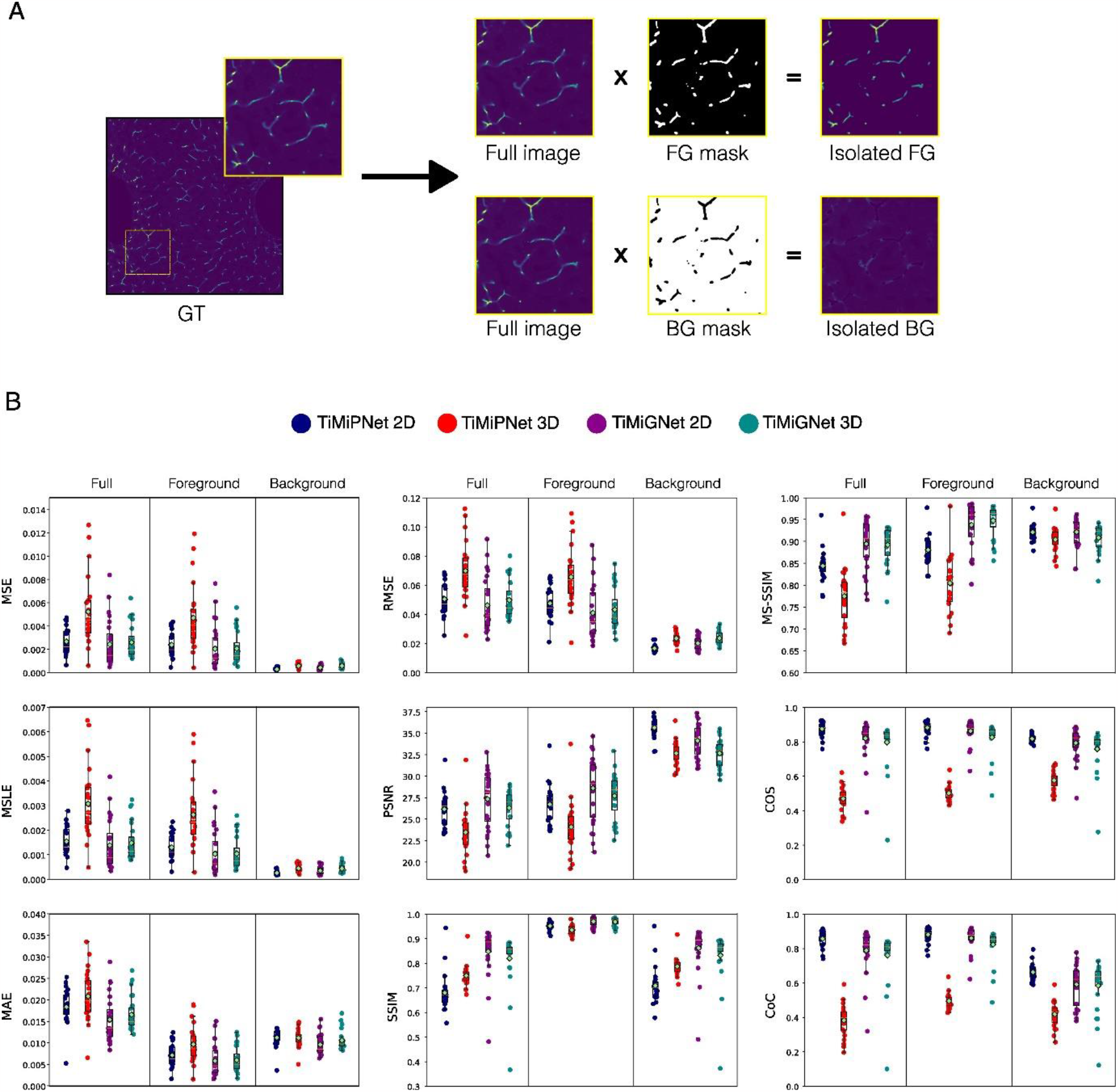
Quantification of the performance of the predictions. a) Schematic representation of the process for generating the mask to split background from foreground in ground truth samples. The masks were calculated using the Otsu method for binary thresholding. b) Quantification of the performance of the predictions of the BC network generated by the 2D and 3D, TiMiPNet, and TiMiGNet models using the following metrics Mean Squared Error, Mean Squared Logarithmic Error, Mean Absolute Error, Root Mean Squared Error, Peak Signal-to-Noise Ratio, Structural Similarity Index Measure, Multi-scale Structural Similarity Index Measure, Cosine Similarity, and Coefficient of Correlation. The test images were splitted in 128x128x128 cubes and the metrics were estimated independently for each cube, i.e. each dot represents one image cube.

**Supp Fig. 2:**
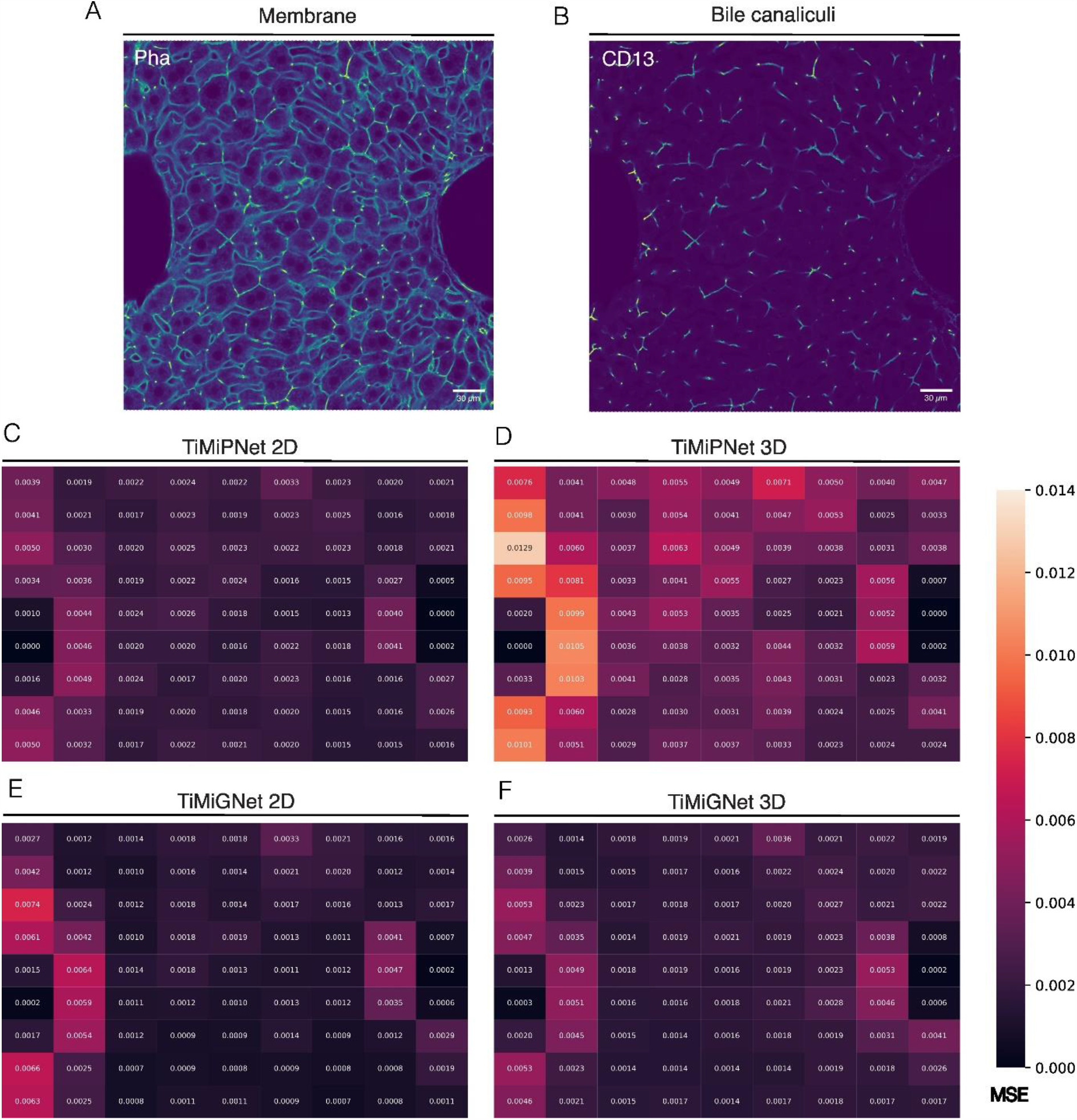
Artificial neural networks show errors close to veins in mouse tissue. (a-b) Representative images actin mesh (membranes), Bile Canaliculi. (c-f) the images were divided into 9x9 blocks and the mean squared error of the predictions of the different models was calculated and shown as a heat map. Whereas low values show good agreement with the ground truth, high vales show potential mispredictions.

## Supplementary Movie Legends

**Supp Movie 1. Z-stack visualization of predicted BC**.

Animation of 2D sections along the axial axis of 3D fluorescent images of the actin mesh (membranes) and Bile Canaliculi (experimental images) and the corresponding predictions of TiMiPNet and TiMiGNet

**Supp Movie 2. 3D rendering of predicted BC**.

3D rendering of the experimental an predicted images of mouse BC.

**Supp Movie 3. Z-stack visualization of predicted sinusoids**.

Animation of 2D sections along the axial axis of 3D fluorescent images of the actin mesh (membranes) and Sinusoids (experimental images) and the corresponding predictions of TiMiPNet and TiMiGNet

**Supp Movie 4. 3D rendering of predicted sinusoids**.

3D rendering of the experimental an predicted images of mouse Sinusoids.

**Supp Movie 5. Z-stack visualization of predicted KCs**.

Animation of 2D sections along the axial axis of 3D fluorescent images of the actin mesh (membranes) and Kupffer cells (experimental images) and the corresponding predictions of TiMiPNet and TiMiGNet

**Supp Movie 6. 3D rendering of predicted KCs**.

3D rendering of the experimental an predicted images of mouse Kupffer cells.

**Supp Movie 7. Z-stack visualization of TiMiGNet-predicted BC for deep tissue reconstruction**.

Animation of 2D sections along the axial axis of 3D fluorescent images of the actin mesh (membranes) and Bile Canaliculi (experimental images) and the corresponding predictions of TiMiPNet and TiMiGNet for deep tissue imaging

**Supp Movie 8. 3D rendering of TiMiGNet-predicted BC for deep tissue reconstruction**.

3D rendering of the experimental an predicted images of mouse BC for deep tissue imaging.

**Supp Movie 9. Z-stack visualization of TiMiGNet-predicted sinusoids for deep tissue reconstruction**.

Animation of 2D sections along the axial axis of 3D fluorescent images of the actin mesh (membranes) and Sinusoids (experimental images) and the corresponding predictions of TiMiPNet and TiMiGNet for deep tissue imaging

**Supp Movie 10. 3D rendering of TiMiGNet-predicted sinusoids for deep tissue reconstruction**.

3D rendering of the experimental an predicted images of mouse Sinusoids for deep tissue imaging.

**Supp Movie 11. Z-stack visualization of TiMiGNet-predicted BC for human tissue** Animation of 2D sections along the axial axis of 3D fluorescent images of the actin mesh (membranes) and BC (experimental images) and the corresponding predictions of TiMiPNet and TiMiGNet for human liver tissue

**Supp Movie 12. 3D rendering of TiMiGNet-predicted BC for human tissue**

3D rendering of the experimental an predicted images of mouse BC for human tissue.

